# Interplay between intraocular and intracranial pressure effects on the optic nerve head in vivo

**DOI:** 10.1101/2020.12.14.422760

**Authors:** Ziyi Zhu, Susannah Waxman, Bo Wang, Jacob Wallace, Samantha E. Schmitt, Elizabeth Tyler-Kabara, Hiroshi Ishikawa, Joel S. Schuman, Matthew A. Smith, Gadi Wollstein, Ian A. Sigal

**Author notes:** Should be considered co-first authors (Ziyi Zhu, Susannah Waxman). Correspondence: Ian A. Sigal, Ph.D. Laboratory of Ocular Biomechanics, Department of Ophthalmology, University of Pittsburgh, Eye & Ear Institute, 203 Lothrop St. Rm. 930, Pittsburgh, PA, USA. 15213, www.ocularbiomechanics.com. <u>Proprietary Interest</u>: J.S. Schuman receives royalties for intellectual property licensed by Massachusetts Institute of Technology to Zeiss. All other authors: Nothing to disclose. <u>Financial support</u>: National Institutes of Health grants R01EY025011, R01EY013178, R01EY023966, R01EY022928, R01EY028662, T32-EY017271 and P30EY008098; Glaucoma Research Foundation Shaffer Grant; Eye and Ear Foundation of Pittsburgh, PA; Research to Prevent Blindness (Support to the Departments of Ophthalmology at the University of Pittsburgh, and at NYU).

## Abstract

Intracranial pressure (ICP) has been proposed to play an important role in the sensitivity to intraocular pressure (IOP) and susceptibility to glaucoma. However, the in vivo effects of simultaneous, controlled, acute variations in ICP and IOP have not been directly measured. We quantified the deformations of the anterior lamina cribrosa (ALC) and scleral canal at Bruch’s membrane opening (BMO) under acute elevation of IOP and/or ICP.

Four eyes of three monkeys were imaged in vivo with OCT under four pressure conditions: IOP and ICP either at baseline or elevated. The BMO and ALC were reconstructed from manual delineations. From these, we determined canal area at the BMO (BMO area), BMO aspect ratio and planarity, and ALC median depth relative to the BMO plane. To better account for the pressure effects on the imaging, we also measured ALC visibility as a percent of the BMO area. Further, ALC depths were analyzed only in regions where the ALC was visible in all pressure conditions. Bootstrap sampling was used to obtain mean estimates and confidence intervals, which were then used to test for significant effects of IOP and ICP, independently and in interaction.

Response to pressure manipulation was highly individualized between eyes, with significant changes detected in a majority of the parameters. Significant interactions between ICP and IOP occurred in all measures, except ALC visibility. On average, ICP elevation expanded BMO area by 0.17mm^2^ at baseline IOP, and contracted BMO area by 0.02 mm^2^ at high IOP. ICP elevation decreased ALC depth by 10μm at baseline IOP, but increased depth by 7 μm at high IOP. ALC visibility decreased as ICP increased, both at baseline (−10%) and high IOP (−17%). IOP elevation expanded BMO area by 0.04 mm^2^ at baseline ICP, and contracted BMO area by 0.09 mm^2^ at high ICP. On average, IOP elevation caused the ALC to displace 3.3 μm anteriorly at baseline ICP, and 22 μm posteriorly at high ICP. ALC visibility improved as IOP increased, both at baseline (5%) and high ICP (8%).

In summary, changing IOP or ICP significantly deformed both the scleral canal and the lamina of the monkey ONH, regardless of the other pressure level. There were significant interactions between the effects of IOP and those of ICP on LC depth, BMO area, aspect ratio and planarity. On most eyes, elevating both pressures by the same amount did not cancel out the effects. Altogether our results show that ICP affects sensitivity to IOP, and thus that it can potentially also affect susceptibility to glaucoma.

**Research Highlights:**

- In vivo ONH deformations caused by acute, controlled, simultaneous changes in IOP and/or ICP can be directly visualized and measured in the monkey eye using OCT.
- Acute changes of either IOP or ICP significantly deformed both the scleral canal and the lamina cribrosa, regardless of the other pressure level.
- Pressures interacted, meaning that the effects of one pressure depended significantly on the level of the other pressure.
- Elevating both pressures did not cancel out the effects of one of them being elevated.
- Our results show that ICP affects sensitivity to IOP, and thus that it can potentially also affect susceptibility to glaucoma.

## 1. INTRODUCTION

Glaucoma is a progressive and irreversible optic neuropathy and the second-leading cause of vision loss in the world ^1,2^. While the mechanisms of neural tissue loss in glaucoma remain unclear, studies suggest that mechanical insult to the optic nerve head (ONH) contributes to the cascade of events that eventually result in neural tissue damage ^3–8^. Much attention has been given to the mechanical insult associated with elevated intraocular pressure (IOP) ^6,9–11^, but the role of intracranial pressure (ICP), which acts on the ONH from outside the eye, is relatively unexplored. Evidence from epidemiological and animal models suggests that ICP could also have an important influence on the ONH, and that it may be a missing factor needed to understand why subjects vary so widely in their sensitivities to elevated IOP ^12–16^.

The in vivo effects on the ONH of acute variations in ICP remain poorly understood. In particular, it remains unclear whether ICP variations can cause deformations of the lamina cribrosa (LC) or the scleral canal. It is also unclear whether the effects of IOP and ICP are independent or if they interact with each other. It has been suggested that the two pressures might counterbalance ^13,15,17–19^. This implies that ONH deformations caused by an elevated IOP might be removed or “cancelled out” by an elevated ICP. Simulations ^14,20^ and experiments ^21^ suggest that the effects of IOP and ICP on the LC can be substantial and do not balance out. However, to the best of our knowledge, the effects of ICP on the ONH, independently and in conjunction with elevated IOP, have not been directly measured in a primate in vivo. To understand the effects of chronically altered pressures on the ONH in glaucoma, we can first work to understand the biomechanics of the ONH under acute changes in pressures.

Our goal was to test if acute changes in ICP or IOP can affect ONH structure and if these variables interact. Interactions would indicate that to understand the effects of one pressure it is necessary to also understand the effects of the other pressure. Specifically, we aimed to quantify in vivo the deformations of the anterior lamina cribrosa (ALC) and scleral canal at Bruch’s membrane opening (BMO) of monkey eyes under acute, controlled variations of IOP and/or ICP.

## 2. METHODS

We used a previously reported ^21^ in vivo monkey model, rhesus macaque, in which IOP and ICP were acutely controlled, independently and simultaneously. Four eyes of three monkeys (M1, M2, M3L, M3R) were imaged in vivo, with optical coherence tomography (OCT). ONH structures were manually delineated in these images. From each scan, five parameters of interest were measured: BMO area, aspect ratio, and planarity as well as ALC median depth and visibility. The parameters were analyzed to test for significant effects of IOP and ICP, independently and in interaction using a bootstrapping approach. Note that the bootstrapping and statistical analysis were designed to evaluate the reliability in the parameters measured, and in detecting their changes upon IOP and ICP changes. The study was not designed to determine whether the measurements in the four eyes studied are representative of a larger population. The details of each of these steps are provided below.

### Animal Handling

Animal handling, pressure control, and imaging were conducted as described elsewhere ^21^. Animal handling followed National Institute of Health (NIH) Guide for the Care and Use of Laboratory Animals, adhered to the Association of Research in Vision and Ophthalmology (ARVO) statement for the Use of Animals in Ophthalmic and Vision Research, and the protocol was approved by the Institutional Animal Care and Use Committee (IACUC) of the University of Pittsburgh. Before the experiment, a clinical examination was conducted to exclude eyes with gross abnormality. For these experiments, animals were initially sedated with 20 mg/kg ketamine, 1 mg/kg diazepam, and 0.04 mg/kg atropine. They were maintained on 1-3% isoflurane for the remainder of the experiment. Animals were put on a ventilator and given vecuronium bromide, a paralytic, intravenously at 0.04-0.1 mg/kg/hr to reduce eye shifting throughout the experiment. Eyes were scanned while animals were in the prone position, with the head held upright and facing the OCT device. The pupils were dilated using tropicamide ophthalmic solution 0.5% (Bausch & Lomb, Rochester, NY). The corneal surface of each eye was covered with a rigid, gas permeable contact lens (Boston EO, Boston, MA) to preserve corneal hydration and improve image quality. The eyes were kept open using a wire speculum and the corneas were hydrated with saline between scans. The animals’ blood pressures and heart rates were monitored throughout the study.

### Pressure Manipulation

For the pressure manipulation we followed the same general approach described elsewhere^21^. After thorough irrigation of the cannula to remove all air bubbles, IOP was controlled by inserting a 27-gauge needle into the anterior chamber and connecting it to a saline reservoir (**Figure 1a**). ICP was controlled by inserting a lumbar drain catheter (Medtronic, Minneapolis, MN) 2.5 cm into the lateral ventricle of the brain and connecting it to a separate saline reservoir. IOP and ICP were thus controlled by adjusting the height of the corresponding reservoir. ICP was also monitored by an ICP pressure also placed into the brain, at least 5mm from the catheter (ICP Express monitoring system, DePuy Synthes, Raynham, MA). Before using the pressure transducer, it was calibrated while submerged in saline solution. IOP and ICP values were controlled within 1 mmHg. Target ICPs were 10, 25, 40 and 5 mmHg. IOPs were set to 15, 30, 50 and 5 mmHg. Based on our experience and the literature, baseline pressures were defined as an IOP of 15 mmHg and an ICP of 10 mmHg ^22,23^. Four pressure conditions were included in this study, one baseline and three experimental with one or both pressures elevated: (IOP/ICP); 15/10 mmHg, 15/25 mmHg, 30/10 mmHg, and 30/25 mmHg.

**Figure 1:**
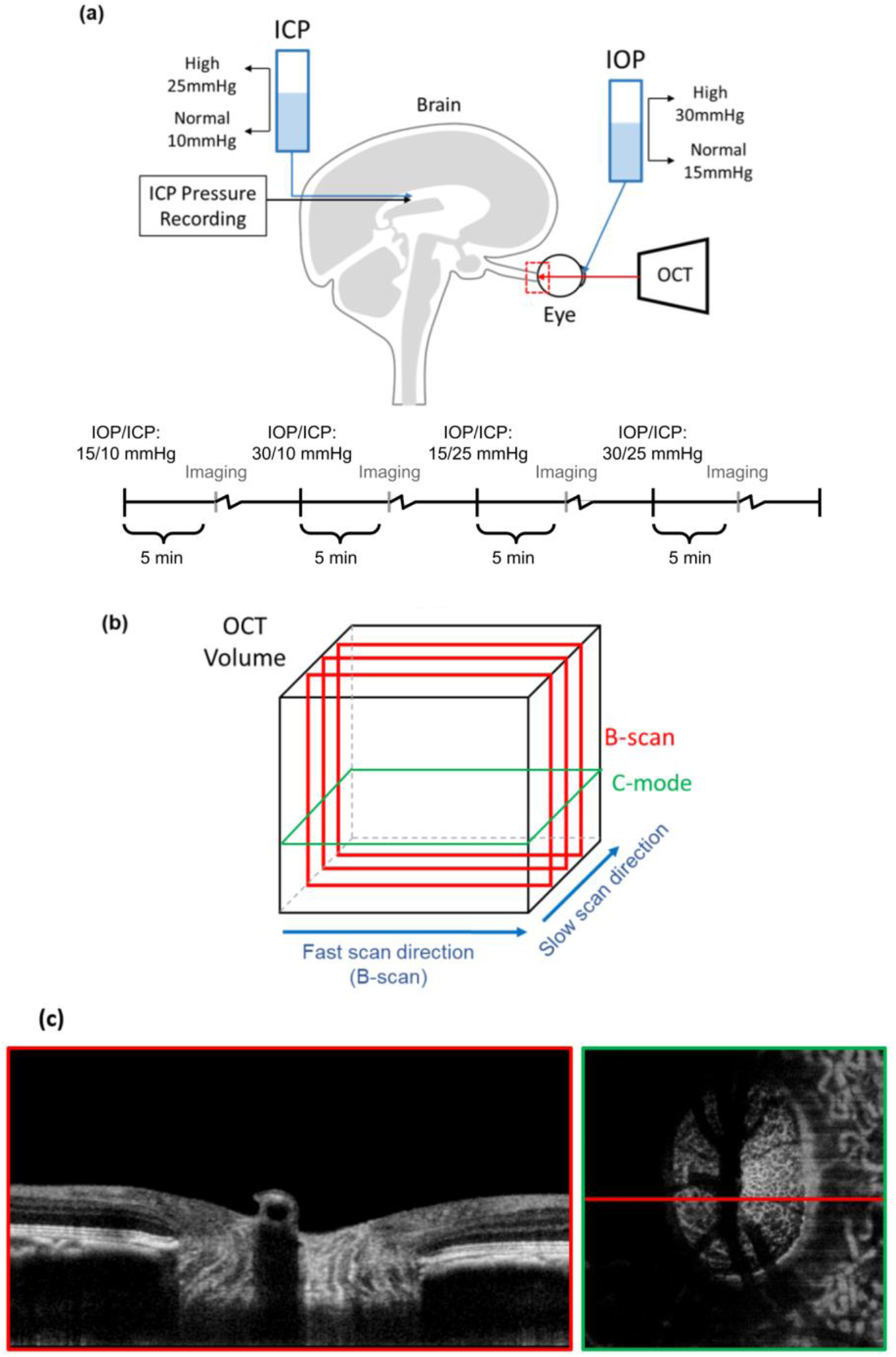
(a) Diagram of in vivo experimental set up and timeline, in which both intraocular and intracranial pressures were controlled via gravity perfusion while the optic nerve head region (red) was imaged with optical coherence tomography. This study focuses on analysis of the four IOP/ICP conditions highlighted. The experimental protocol included other IOP/ICP conditions between the ones highlighted. See the main text for details. Motion artifacts in the slow scan direction were removed (b). Example B-scan and C-mode views at the lamina cribrosa level acquired with an IOP of 15 mmHg and ICP of 8 mmHg (c).

### Imaging

Eyes were imaged with spectral domain optical coherence tomography (OCT) (Bioptigen, Research Triangle, NC) with a scan rate of 20,000 A-scans/second, modified with a broadband superluminescent diode (Superlum, Dublin, Ireland, λ = 870 nm, Δλ = 200 nm). OCT volume scans were acquired with a 5mm × 5mm × 2mm (512 × 512 × 1024 pixels sampling, with 1 Frame per B-scan, i.e. no repetitions) setting, centered on the ONH region (**Figure 1a**). Multiple scans were obtained at each pressure condition, and the best quality scan at each pressure condition was chosen for manual delineation. Image quality criteria are detailed elsewhere ^21^. Image quality tended to decrease with increasing anesthesia time. To ensure that image quality remained high and to minimize the amount of time each animal spent under anesthesia, in the early experiments we imaged only one eye. After becoming more comfortable and quicker with our animal protocols, imaging could be performed fast enough to capture images from both eyes in the third monkey. After each IOP and/or ICP change we stepped back for 5 minutes, waiting for the eyes to stabilize. In addition, at each pressure we spent 20-30 minutes adjusting equipment and conducting the imaging. Exact times varied slightly depending on how quickly we were able to get the imaging setup, keeping the cornea hydrated, etc. Since only a subset of images were analyzed for this work, the actual times between pressures were always at least 60min for Monkeys 1 and 2, and 45 min for Monkey 3. Based on our experience and the literature, we believe that these times are sufficient to minimize viscoelastic effects.^24–27^.

All scans were re-sampled at 1 × 1 × 1 scale for analysis ^28^ Eyes vary in optical power and OCT systems are optimized for imaging human eyes. Hence, OCT images of monkey ONHs must be rescaled in the transverse dimensions. To set the dimensions, we followed the process described previously ^21^. Briefly, after the experiment, eyes were enucleated, processed for histology, and sections were imaged with polarized light microscopy. The images were reconstructed into 3D stacks and used to obtain eye-specific transverse scaling factors based on the dimensions of the scleral canal at BMO. Elsewhere we have shown that histological processing does not alter the scale of eye tissues ^29^.

### Delineation

Delineations were done by an experienced observer, masked to both IOP and ICP conditions, in an open-source imaging processing package, Fiji ^30^. Two ONH landmarks, the scleral canal at BMO and the anterior boundary of the LC (ALC), were delineated in equally spaced OCT B-scans (**Figure 1**). The ALC was sampled at a higher transverse resolution, every 31 μm, than the BMO, every 62 μm, to best resolve its comparatively non-uniform structure. The BMO best-fit plane was used as a reference for measurements of BMO planarity and ALC depth. The ALC and scleral canal at BMO were selected for analysis because they have often been used in studies of monkey ONH biomechanics ^31^, and because simulation analyses have shown that they are useful to capture essential elements of ONH biomechanics ^32^.

### 3D Reconstruction & Registration

Motion artifacts, from breathing, heartbeat, or surgical table vibrations were discernible in the slow-scan (superior-inferior) direction as a wavy pattern in the otherwise smooth structure of Bruch’s membrane. These were removed by translating B-scan images in the anterior-posterior direction. Custom scripts were used to import manual delineations of BMOs made on virtual superior-inferior volume cross-sections from Fiji into Matlab (Mathworks, Natick, MA, USA). Custom scripts were also used to interpolate between the scattered manual markings of the BMO and ALC. This allowed us to obtain 3D reconstructions for analysis (**Figure 2**). When mitigating motion artifacts, we used positions as far as possible from the canal at BMO as landmarks for alignment. This was done to minimize alignment-based changes impacted by changes of BMO planarity themselves. Additionally, we filtered motion artifacts with frequencies corresponding to heartbeat. Images of the ALC across different pressure conditions within an eye were registered by aligning the center and principal axes of the BMO best-fit plane.

**Figure 2:**
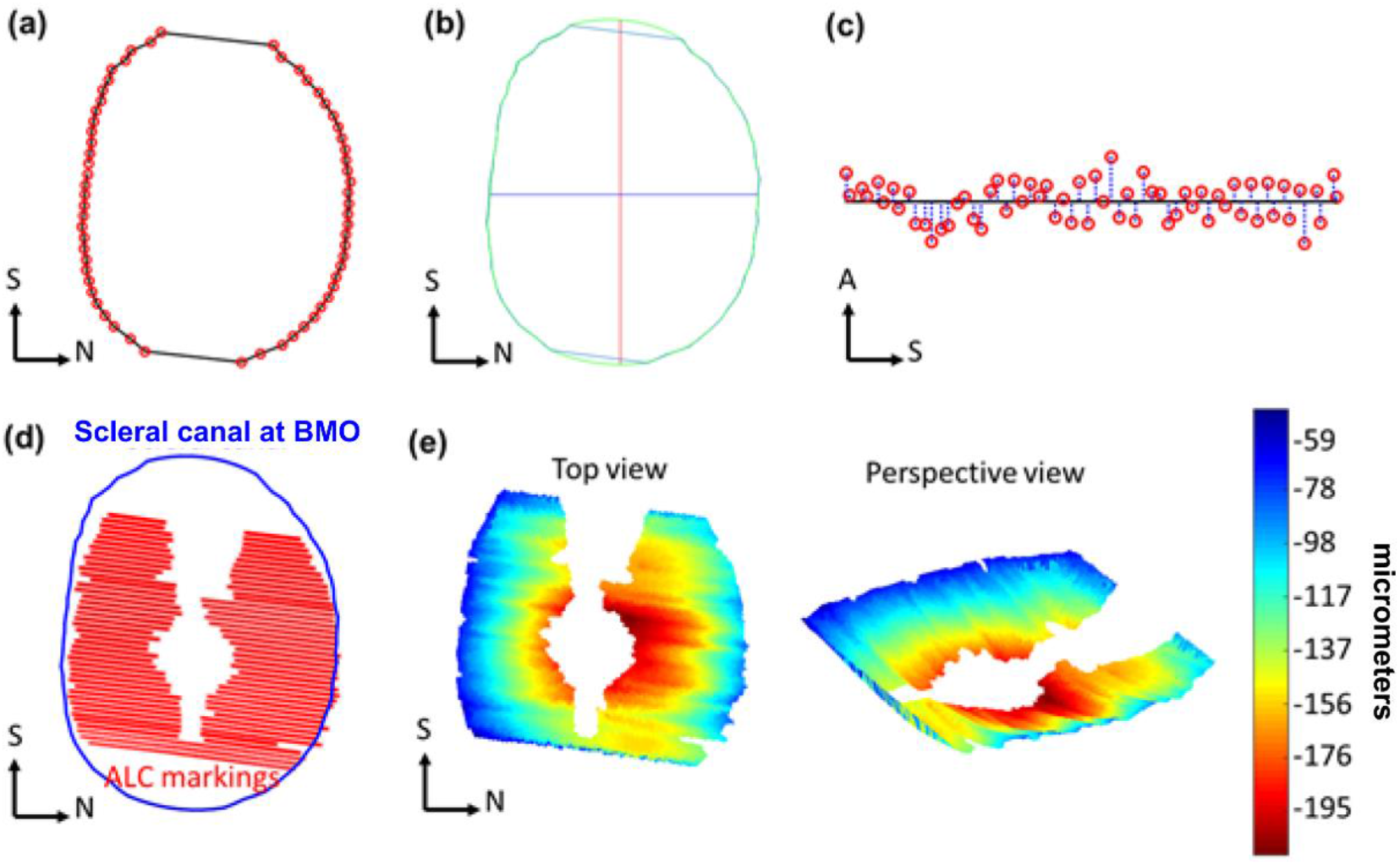
Example markings of Bruch’s membrane opening, (BMO, red) (a). Example scleral canal area (within green perimeter), interpolated from BMO markings, and its corresponding principal axes (red, blue) (b). Scleral canal planarity was calculated as the average of distances (blue) from BMO markings (red) to BMO best-fit plane (black) (c). Example scleral canal (blue) and anterior lamina cribrosa (ALC) markings (red) used to reconstruct ALC surface and compute ALC depth (d). Heat maps of ALC depth (shallow to deep: blue to red) (e). S: Superior, N: Nasal, A: Anterior.

### BMO Area, Aspect Ratio, Planarity

BMO area was computed as the projected area of BMO on the best-fit plane. BMO aspect ratio was computed from the ratio between the major and minor principal axes of this plane (**Figure 2a-c**). The planarity of the BMO was defined as the average of the distances from BMO points to the best-fit plane, measuring the extent to which the BMO deviated from a flat plane ^8,33^. Note that with this definition, a perfectly planar canal opening has a planarity of zero, with planarity increasing as the shape deviates from the perfect plane.

### Lamina-Visibility, Median Depth

The 3D ALC surface was projected onto the BMO best-fit plane and ALC visibility, or analyzable ALC, was computed as percent of projected ALC area normalized to baseline BMO area (**Figure 2e-d**). To avoid potential biases due to variable LC visibility with IOP, ALC median depth, relative to BMO best-fit plane, was measured in regions where ALC was visible in all pressure conditions within each eye. We defined the sign of the depth with positive direction being anterior to the BMO and vice versa.

### Repeatability

Repeatability of measurements was evaluated as we have done previously ^34^. Briefly, an OCT volume was processed and marked three times for each of five parameters and standard deviations over the three markings used as a measure of repeatability.

### Bootstrap Sampling

Bootstrap sampling was used to assess the reliability of observed structural deformations for each eye and make the best possible use of the limited number of monkeys and eyes. A custom Matlab script was used to randomly select a subset of 75% of the B-scan markings in each volume, for each ONH. These sampled markings were then used to reconstruct and compute the five canal and ALC parameters, as described above. The procedure was repeated 10 times to obtain sampling distributions for each parameter within each eye at each pressure condition. **Figure 3** illustrates the process of bootstrap sampling and subsequent statistical analyses.

**Figure 3:**
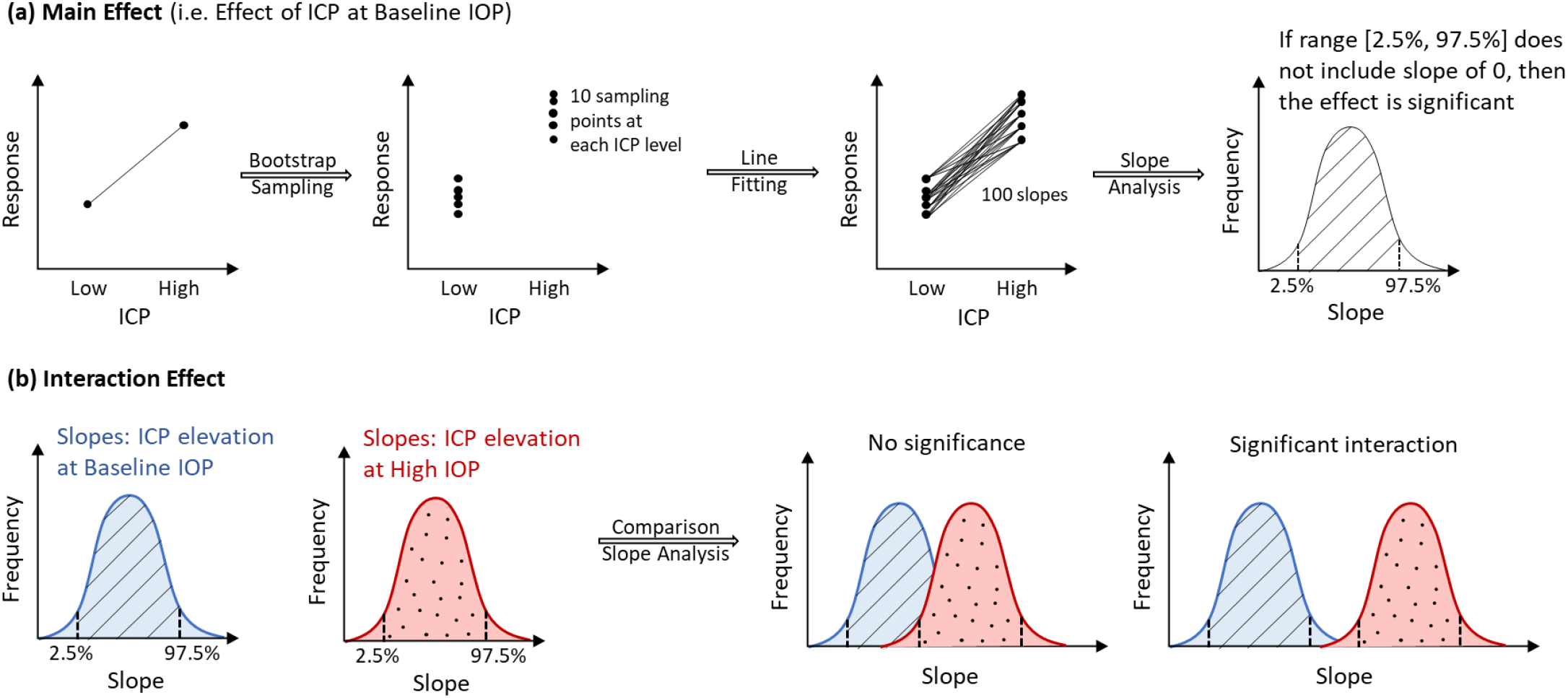
Diagram of statistical tests for the main effects of IOP, ICP (a) and the interaction of their effects (b). (a) Main effect: Bootstrap sampling was used to generate 20 sampling points, 10 sampling points at each of 2 ICP levels. Fitting lines through these points, 100 slopes and their 95% range (between 2.5% and 97.5%), were computed. A significant main effect was detected if the range did not include a slope of 0. (b) Interaction effect: From left to right: Similar procedure in (a) is performed to generate slopes and their two corresponding ranges due to ICP elevation at baseline and at high IOP. A significant interaction between ICP effects and IOP effects was detected if these two ranges did overlap.

### Estimates & Percentage changes

Within each eye, the estimates of each ONH parameter at each pressure condition were computed as the mean of the bootstrap distribution. The confidence intervals (CI) of the estimates were defined as the 95^th^ percentiles (between the 2.5% and 97.5% percentiles). To compute these percentiles, the bootstrap distributions were fit by a normal distribution and the percentiles estimated from the normal. A small CI indicates that the bootstrap distribution is tight, indicating that selecting different subsets of the markings leads to similar measures. Conversely, a wide CI indicates that the bootstrap distribution is wide, indicating that the measure is sensitive to the specific set of markings from which it was derived.

The percentage change of the estimate at each pressure condition was calculated with respect to the baseline estimate. Positive percentage changes of ALC median depth corresponded to more anterior ALCs (and thus, negative changes to more posterior ALCs), and positive percentage changes of BMO area corresponded to scleral canal expansions (and thus negative changes to scleral canal contractions).

### Statistical analysis

For each parameter in each eye, we computed the statistical significance of the independent (main effect) and interaction effects of IOP and ICP as described in **Figure 3**. A main effect was deemed significant when the range of slopes within the 95% confidence interval did not include 0. An interaction effect was deemed significant if the ranges of slopes within the 95% confidence interval did not overlap. Mean changes of significantly different cases were computed and reported.

## 3. RESULTS

We successfully acquired images and measurements on all eyes and conditions, except for two cases. No eye of the monkeys reported here had a gross abnormality. Images of the left eye of Monkey 3 at elevated IOP, at both baseline and elevated ICP, did not have good LC visibility and were therefore removed from analysis. When measuring repeatability, standard deviations over the three markings were 0.01, 0.01 mm^2^, 0.44 μm for BMO aspect ratio, area and planarity, respectively. For ALC depth and visibility, the standard deviations were 5.5 μm and 1%, respectively. For comparison, the voxel edge length of the OCT images was 3.125 μm.

Deformations of ONH structures resulting from IOP and ICP variations were evident by overlaying delineations in corresponding B-scans (**Figure 4**). Baseline parameters are summarized in **Table 1**. The scleral canal at BMO was evaluated as an indicator of greater scleral canal morphology. The effects of ICP elevation on the shape of the scleral canal at BMO, at high and low IOP are illustrated in Figure 5. Overall, pressure changes had significant effects on the various morphological parameters. The effects are shown in **Figures 6** to **8** and summarized in **Tables 2** and **3**. Results of the statistical tests are summarized in **Figure 9**. The location and direction of force generated by IOP and ICP are illustrated in **Figure 10**.

**Figure 4:**
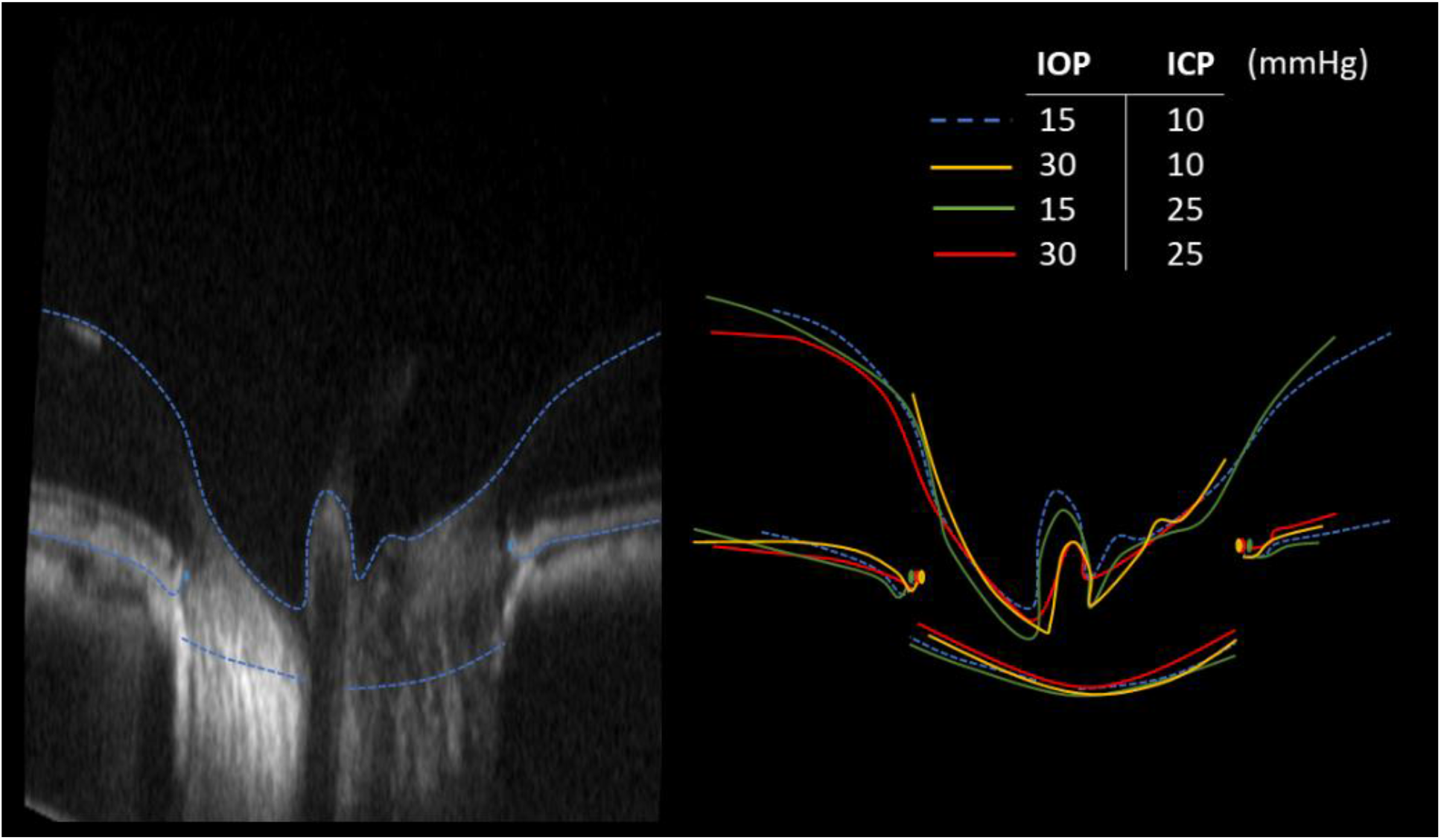
Example qualitative comparison of effects of IOP and ICP. (Left) Baseline B-scan and markings. (Right) Overlay markings from all pressure conditions on baseline B-scan to demonstrate deformations of ONH structures. For this study we analyzed the scleral canal at BMO and the anterior boundary of the LC. In this image we also show delineations of the BM and the inner limiting membrane (including over the central retinal vessels). The dashed lines are delineations at the baseline IOP and ICP levels. Note that to simplify discerning the differences, the B-scan and outlines are shown exaggerated 3 times in axial direction, as is the common for presenting OCT.

**Figure 5:**
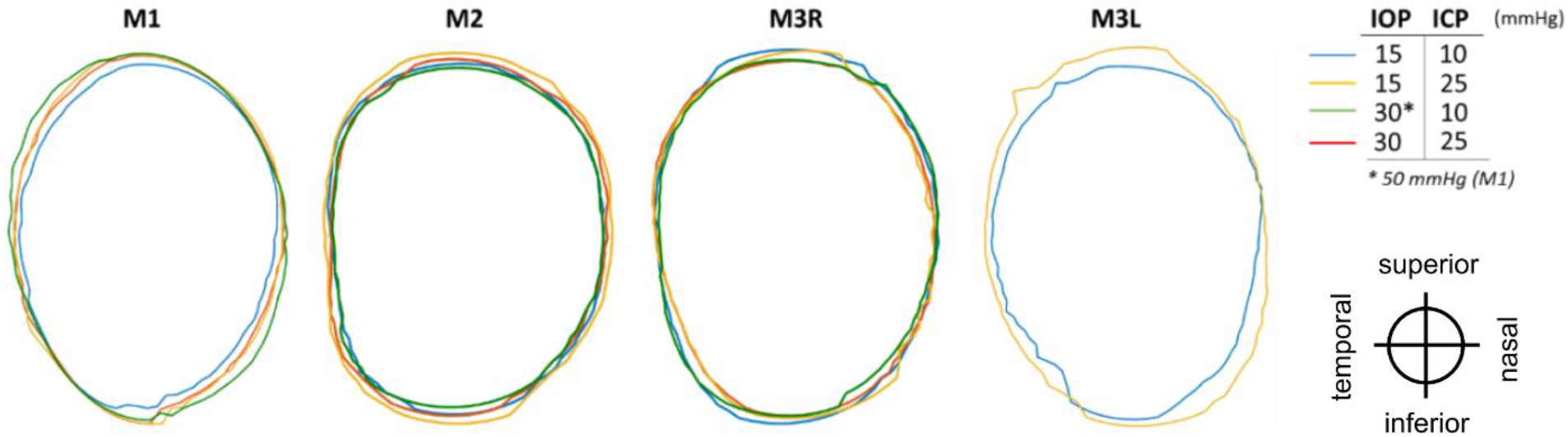
Outlines of the scleral canal at the Bruch’s membrane openings, for each eye at 4 pressure conditions: baseline (blue), base IOP/high ICP (yellow), high IOP/base ICP (green), and high IOP/high ICP (red). Outlines were registered rigidly by the centroid and principal axes. Images of M3L at elevated IOP had poor LC visibility and were therefore excluded from analysis. Orientation of eyes as displayed is indicated at the lower right-hand side.

**Figure 6:**
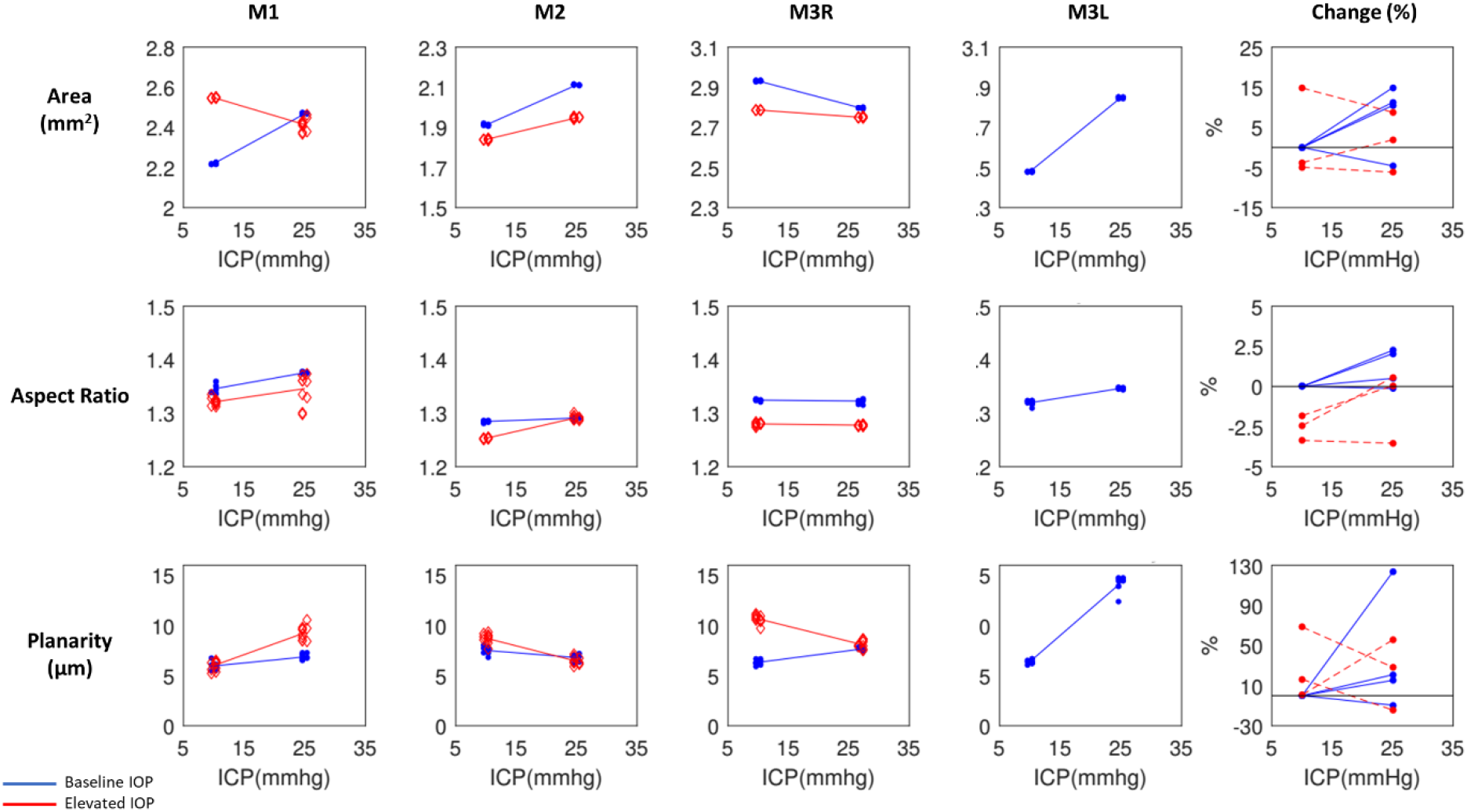
Scleral canal displacements due to variations in intraocular (IOP) and intracranial (ICP) pressures. Percentage changes of BMO area, aspect ratio, and planarity with respect to baseline values due to ICP elevation at baseline IOP (blue) and elevated IOP (red). Each line represents the regression of the estimates, or average of 10 bootstrap sampling points, at each ICP. To reduce overlap, the symbols were scattered laterally slightly.

**Figure 7:**
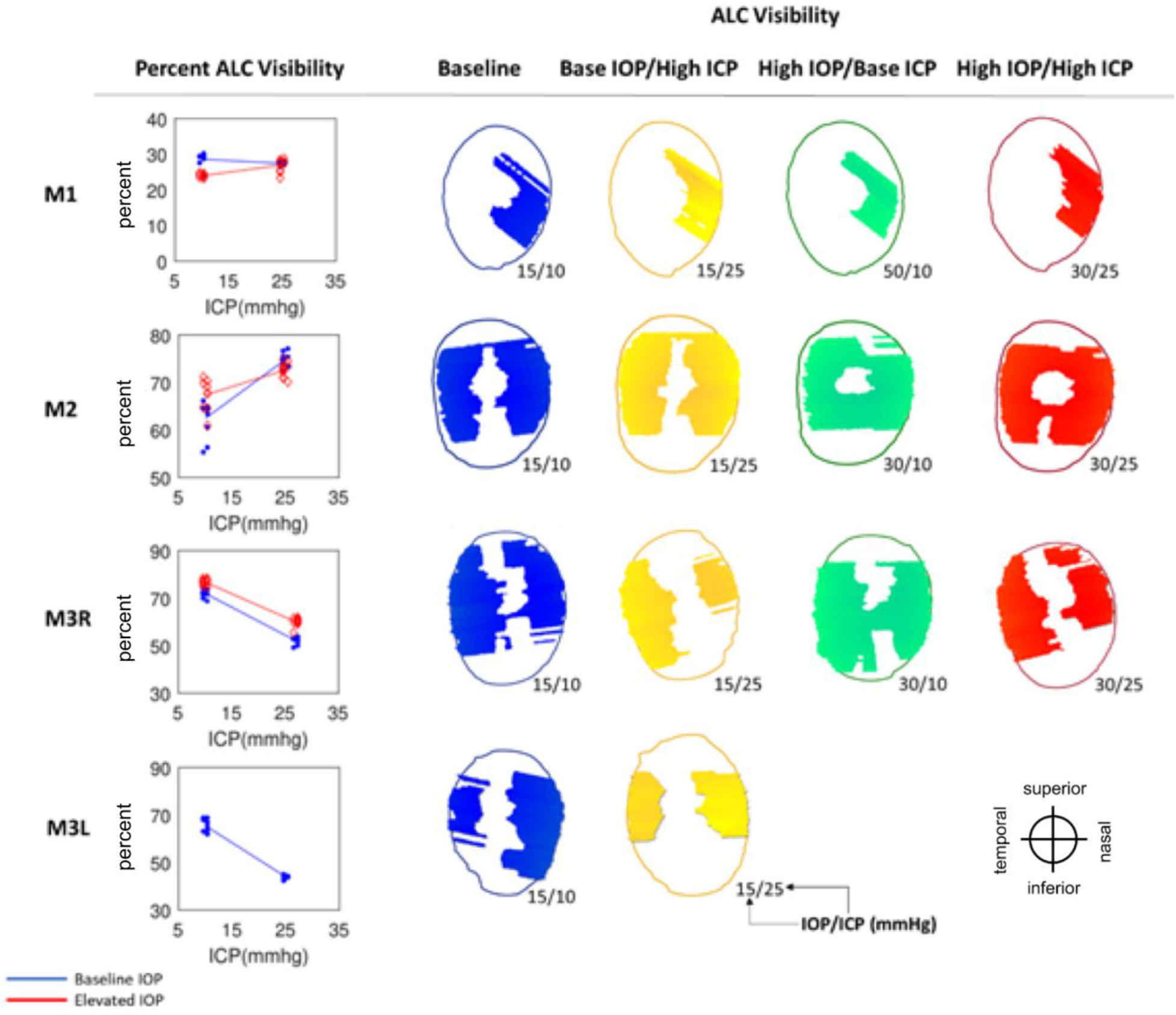
Anterior lamina cribrosa (ALC) visibility. (Left) Percentage of ALC visibility at baseline IOP (blue) and elevated IOP (red). Each line represents the regression of the estimates, or average of 10 bootstrap sampling points, at each ICP. (Right) Maps of ALC visibility. Shown are canal outline (thin line) and ALC for 4 pressure conditions: baseline (blue), baseline IOP/high ICP (yellow), high IOP/baseline ICP (green), and high IOP/high ICP (red). Pressures for each condition are indicated as IOP/ICP.

**Figure 8:**
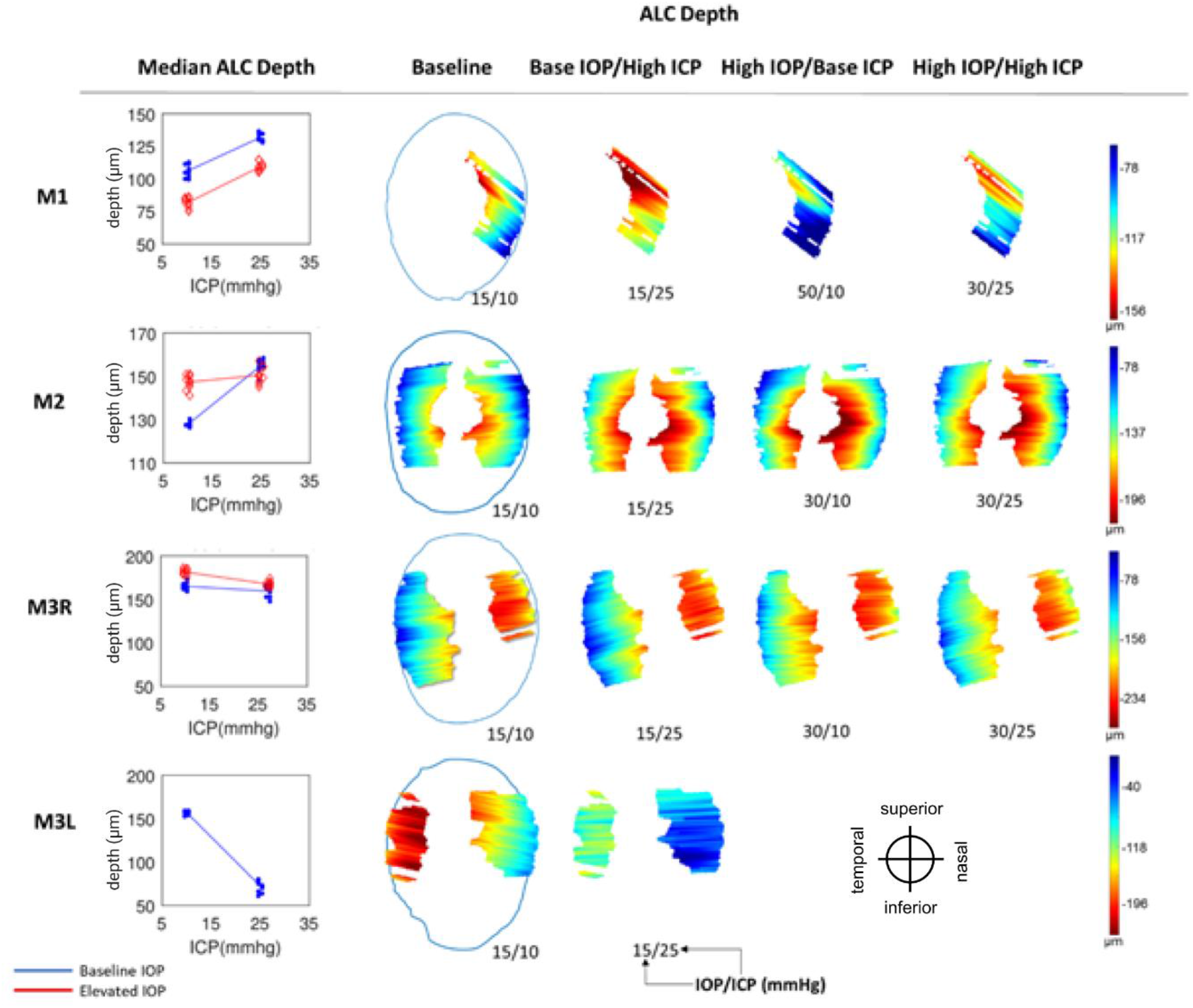
Anterior lamina cribrosa (ALC) depth. (Left) Median ALC depth at baseline IOP (blue) and elevated IOP (red). Each line represents the regression of the estimates, or average of 10 bootstrap sampling points, at each ICP. (Right) Heat maps of ALC depth (blue to red: shallower to deeper) with respect to scleral canal (blue outline), shown only on regions visible across all 4 pressure conditions within an eye. Pressures for each condition are indicated as IOP/ICP.

**Figure 9:**
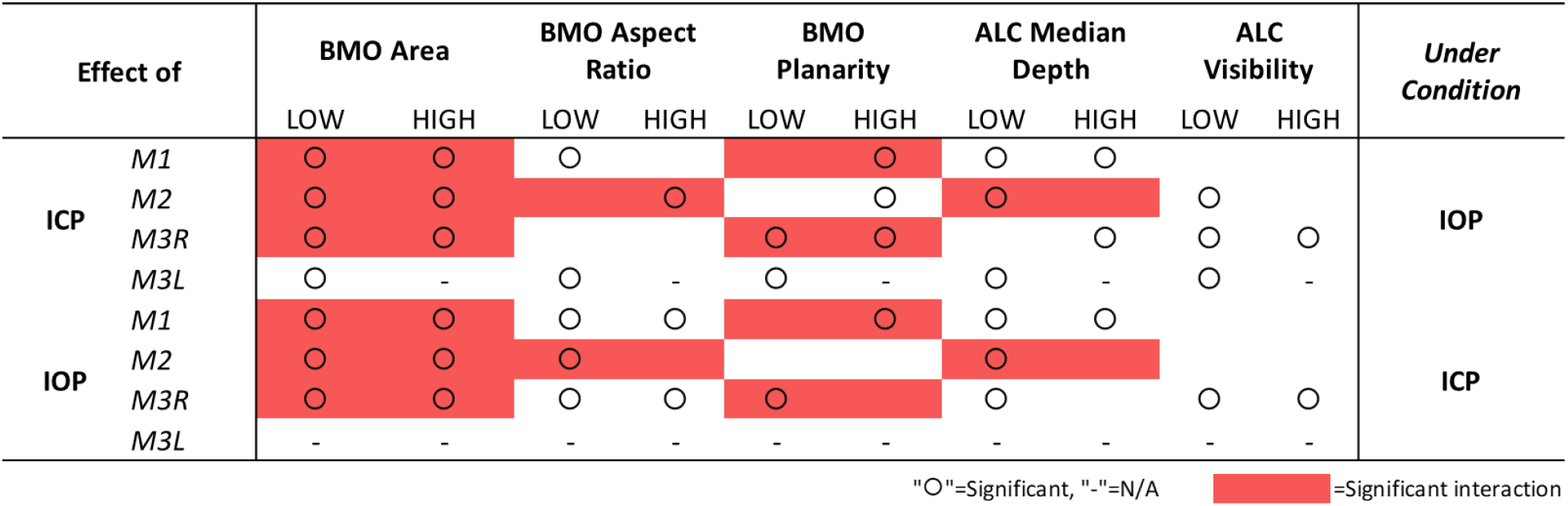
Summary of statistical results showing the significance of IOP and ICP independent effects (“O”) as well as the significance of their interaction effects (red box) on ONH structures. Scleral canal measurements taken at BMO.

**Figure 10:**
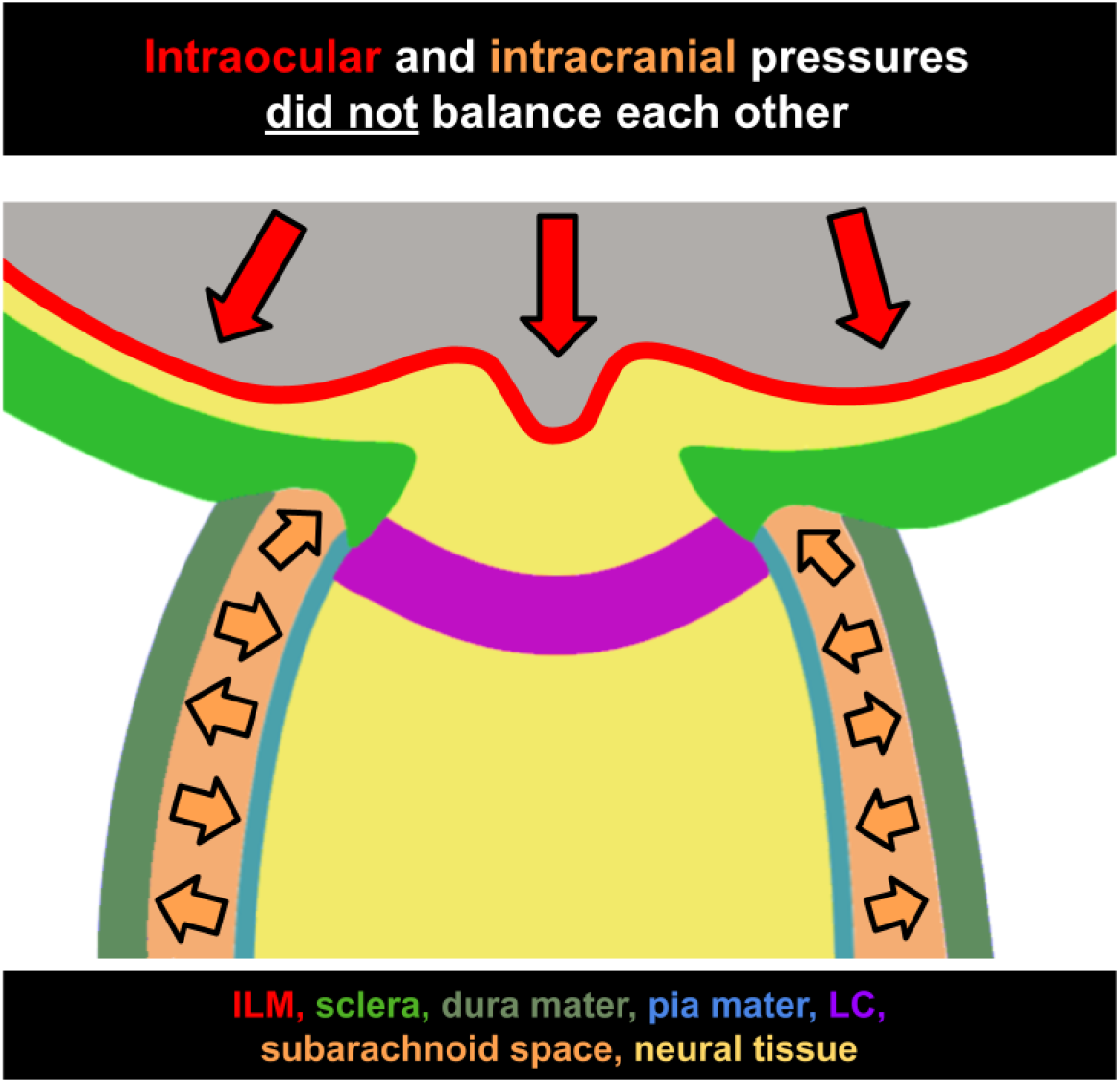
Intraocular and intracranial pressures do not balance each other. Diagram of the ONH with representations of the inner limiting membrane (ILM, red), sclera (green), dura mater (gray-green), pia mater (blue), lamina cribrosa (LC, purple), subarachnoid space (orange), and neural tissue (yellow). Direction of force placed by intraocular pressure (red arrows) and intracranial pressure (orange arrows.)

**Table 1:**
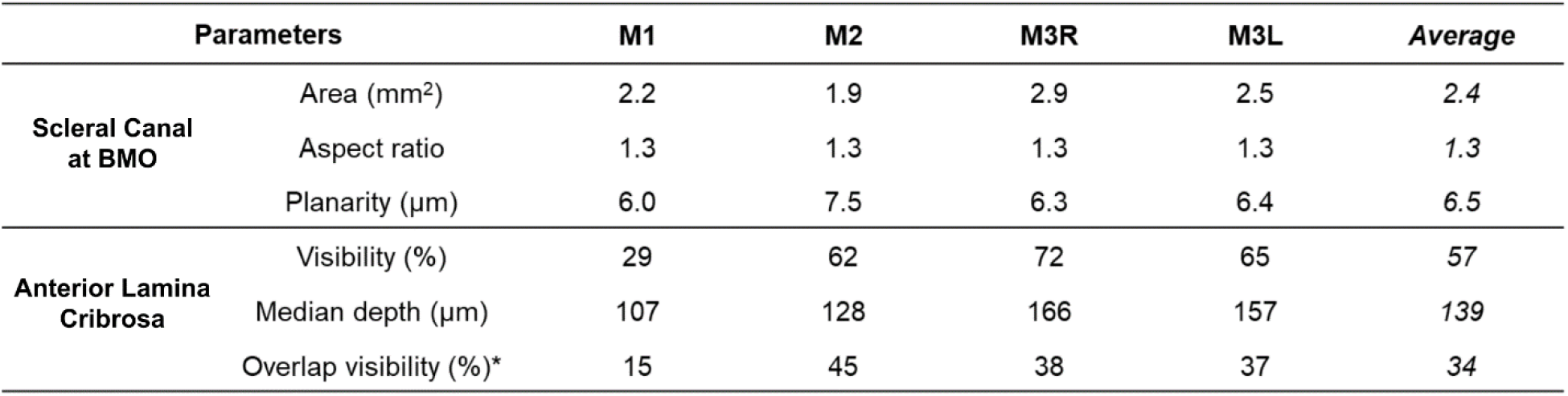
Baseline parameters of the scleral canal, measured at the Bruch’s membrane opening, and of the anterior lamina cribrosa across 4 eyes. Overlap visibility (*) is the common visible region of the lamina across all analyzed conditions within each eye.

| Parameters |  | M1 | M2 | M3R | M3L | Average |
| --- | --- | --- | --- | --- | --- | --- |
| Scleral Canal at BMO | Area (mm<sup>2</sup>) | 2.2 | 1.9 | 2.9 | 2.5 | 2.4 |
|  | Aspect ratio | 1.3 | 1.3 | 1.3 | 1.3 | 1.3 |
|  | Planarity (μm) | 6.0 | 7.5 | 6.3 | 6.4 | 6.5 |
| Anterior Lamina Cribrosa | Visibility (%) | 29 | 62 | 72 | 65 | 57 |
|  | Median depth (μm) | 107 | 128 | 166 | 157 | 139 |
|  | Overlap visibility (%)* | 15 | 45 | 38 | 37 | 34 |

**Table 2:** Summary of changes of ONH parameters as a response to pressure variations. LOW: baseline pressure; HIGH: elevated pressure. These ranges of the parameters were used to test the significance of parameter effects and interactions. Dashes indicate relationships that could not be computed due to images of insufficient quality.

| Effect of |  | BMO Area (mm<sup>2</sup>) |  | BMO Aspect Ratio |  | BMO Planarity (μm) |  | ALC Median Depth (μm) |  | ALC Visibility (%) |  |
| --- | --- | --- | --- | --- | --- | --- | --- | --- | --- | --- | --- |
|  |  | LOW | HIGH | LOW | HIGH | LOW | HIGH | LOW | HIGH | LOW | HIGH |
| ICP | M1 | 0.25 | -0.13 | 0.030 | 0.025 | 0.9 | 3.3 | 25 | 28 | -1 | 3 |
|  | M2 | 0.20 | 0.11 | 0.006 | 0.039 | -0.7 | -2.3 | 28 | 3 | 12 | 5 |
|  | M3R | -0.14 | -0.04 | -0.002 | -0.002 | 1.3 | -2.5 | -8 | -14 | -20 | -17 |
|  | M3L | 0.37 | - | 0.027 | - | 7.9 | - | -83 | - | -22 | - |
| IOP | M1 | 0.33 | -0.05 | -0.024 | -0.030 | 0.0 | 2.4 | -25 | -22 | -5 | 0 |
|  | M2 | -0.07 | -0.16 | -0.031 | 0.001 | 1.2 | -0.4 | 20 | -5 | 5 | -2 |
|  | M3R | -0.14 | -0.05 | -0.045 | -0.045 | 4.3 | 0.5 | 15 | 9 | 5 | 8 |
|  | M3L | - | - | - | - | - | - | - | - | - | - |

**Table 3:** Summary of mean regression slopes corresponding with cases which demonstrated significant effects of IOP, ICP, and IOP-ICP interaction. LOW: baseline pressure; HIGH: elevated pressure. Dashes indicate relationships that could not be computed due to images of insufficient quality. Blank cells indicate results that were not significant.

| Effect of |  | BMO Area (mm<sup>2</sup>) |  | BMO Aspect Ratio |  | BMO Planarity (μm) |  | ALC Median Depth (μm) |  | ALC Visibility (%) |  |
| --- | --- | --- | --- | --- | --- | --- | --- | --- | --- | --- | --- |
|  |  | LOW | HIGH | LOW | HIGH | LOW | HIGH | LOW | HIGH | LOW | HIGH |
| ICP | M1 | 0.017 | -0.009 | 0.002 |  |  | 0.217 | 1.736 | 1.857 |  |  |
|  | M2 | 0.013 | 0.007 |  | 0.003 |  | -0.155 | 1.821 |  | 0.818 |  |
|  | M3R | -0.008 | -0.002 |  |  | 0.078 | -0.170 |  | -0.927 | -1.173 | -1.114 |
|  | M3L | 0.025 | - | 0.002 | - | 0.527 | - | -5.650 | - | -1.435 | - |
| IOP | M1 | 0.009 | -0.004 | -0.001 | -0.002 |  | 0.159 | -0.693 | -1.497 |  |  |
|  | M2 | -0.005 | -0.011 | -0.002 |  |  |  | 1.304 |  |  |  |
|  | M3R | -0.010 | -0.002 | -0.003 | -0.003 | 0.290 |  | 1.087 |  | 0.314 | 0.529 |
|  | M3L | - | - | - | - | - | - | - | - | - | - |

On average, ICP elevation expanded canal area at BMO by 0.17mm^2^ at baseline IOP and contracted the BMO area by 0.02 mm^2^ at high IOP. ICP elevation decreased ALC depth by 10μm at baseline IOP, but increased depth by 7μm at high IOP. ALC visibility decreased as ICP increased, both at baseline (−10%) and high IOP (−17%). IOP elevation expanded BMO area by 0.04 mm^2^ at baseline ICP, and contracted BMO area by 0.09 mm^2^ at high ICP. On average, IOP elevation caused the ALC to displace 3.3 μm anteriorly at baseline ICP, and 22 μm posteriorly at high ICP. ALC visibility improved as IOP increased, both at baseline (5%) and high ICP (8%).

### BMO Area, Aspect Ratio, Planarity

For BMO area, ICP elevation had a significant effect at both high and low IOP on 3 eyes (M1-M3R) and at low IOP on M3L (**Figures 5** and **6**, **Tables 2** and **3**). IOP elevation had significant effects at both high and low ICP on 3 eyes (M1-M3R). There were significant interactions of ICP and IOP effects on BMO area in 3 eyes (M1-M3R).

For BMO aspect ratio, ICP elevation had significant effects at low IOP on 2 eyes (M1, M3L) and at high IOP on M2 (**Figure 6**, **Tables 2** and **3**). IOP elevation had significant effects at both high and low ICP on 2 eyes (M1, M3R) and at low ICP on M2. There was 1 significant interaction of ICP and IOP effects on BMO aspect ratio in M2.

For BMO planarity, ICP elevation had significant effects at low IOP on 2 eyes (M3R, M3L) and at high IOP on 3 eyes (M1-M3R) (**Figure 6**, **Tables 2** and **3**). IOP elevation had significant effects at low ICP on M3R and at high ICP on M1. There were significant interactions of ICP and IOP effects on BMO planarity in 2 eyes (M1, M3R).

Considering all cases, pressure variations induced the largest changes on BMO planarity (−14% to 123%), followed by the BMO area (−6% to 15%), and finally by the BMO aspect ratio with very small changes (−4% to 2%) (**Figure 6**).

### Lamina Depth and Visibility

For ALC median depth, ICP elevation had significant effects at low IOP on 3 eyes (M1, M2, M3L) and at high IOP on 2 eyes (M1, M3R) (**Figure 7**, **Tables 2** and **3**). IOP elevation had significant effects at low ICP on 3 eyes (M1-M3R) and at high ICP on M1. The effects of IOP and ICP on ALC depth had a significant interaction for M2.

Across the 4 eyes, baseline ALC visibility and median depth average were 57% and 139μm, respectively (**Table 1**). The amount of ALC visible in scans at each pressure condition varied between eyes, ranging from 15-45%, with an average of 34%. Regardless of pressure conditions, ALC visibilities were higher in M2, M3R, and M3L. These monkeys had analyzable ALC in both nasal and temporal sides, compared to M1L, in which ALC was visible only on the temporal side (**Table 1**, **Figure 7**). ICP elevation had significant effects on ALC visibility at low IOP and at high IOP (**Figure 7**, **Tables 2** and **3**). No significant interaction between ICP and IOP effects on ALC visibility was detected.

Considering all pressure settings across the 4 eyes, changes in both depth and visibility of the LC were substantial. Median depth changed between −53% and 24% and visibility changed between −33% and 20% (**Supplemental Figure 1**).

### Heterogeneity of Responses to IOP/ICP

The variable responses in structural parameters are best visualized in **Figures 5–9** and **Tables 2** and **3**. To help readers interpret the figures and understand their implications, we will consider some examples. For instance, consider the ALC Median depth of M1 in **Figure 8**. As ICP was increased at baseline IOP (blue line) we measured an increase in the depth of the ALC. All blue points representing the data subsets created for bootstrapping were neatly clustered around each median depth, indicating a relative homogeneity of depth measurements and thus good confidence in the values. A regression line was fit between the clusters at baseline and elevated ICP settings for visualization. The positive, non-zero slope represented a significant increase in ALC depth as a function of ICP. When we repeated the experiment at elevated IOP, we observed a similar relationship, but the ALC depths were shifted shallower at both ICP settings. The parallel lines indicate that ICP and IOP affected the deformations independently and that there was no interaction between variables. This first row of **Figure 8** summarizes a case in which ICP elevation had a significant effect at each IOP, but there was no interaction between ICP and IOP. The following subject, M2, showed a similar relationship for elevations of ICP at baseline IOP (blue line), but at high IOP the relationship was not present (red line). This indicates that the variables interacted strongly. For M3L, ALC depth was reduced with increasing ICP. No significant interaction was observed between the effects of ICP and IOP for ALC depth in M3L. These variable structural response are summarized in **Table 2**, where it can be seen that each column has at least one negative and one positive value. **Figure 9** summarizes the statistical results and the significance of IOP and ICP effects independently and in interaction. These examples are representative of the heterogeneity in the ONH response between eyes that was present for all parameters.

Interestingly, in most cases, setting the translaminar pressure difference (TLPD) to the baseline levels did not deform the canal and lamina back to baseline configurations. For example, cases in which IOP was 15 and ICP was 10 mmHg (baseline for all eyes) can be compared with cases in which IOP was 30 and ICP was 25 mmHg (such as in M1, M2, and M3R). Each of these cases had a TLPD of 5 mmHg. However, we observed changes in scleral canal displacement (**Figure 6**), ALC visibility (**Figure 7**), and ALC depth (**Figure 8**) upon pressures increases despite conserved TLPD. In M1 and M2, scleral canal area, ALC visibility, ALC depth were all increased as IOP and ICP were increased from baseline to elevated pressures. In M3R, these factors were respectively decreased, decreased, and similar to their respective baselines. Based on this data, we did not observe a strong relationship between TLPD and ONH morphology.

Figure 10 shows the location of where forces from IOP and CSFP are applied in effort to illustrate why IOP and ICP do not balance each other out. Provided this context, it is clear why TLPD is not a strong predictor of ONH morphology. IOP and ICP can undergo equal changes in magnitude, maintaining a consistent TLPD, but they cannot be expected to have opposite effects due to the locations at which they are applied.

## 4. DISCUSSION

In this study, we set out to quantify in vivo deformations of the ALC and scleral canal under acute, controlled variations of IOP and/or ICP in a monkey model. Four main findings arise from this work: (1) changes in either IOP or ICP caused significant, detectable ONH deformations, (2) there were strong interactions between the effects of IOP and ICP, (3) elevating both pressures by the same amount, to maintain TLPD, did not cancel out the effects, and (4) a high degree of heterogeneity in response to pressure manipulation was observed among eyes. Our findings are important because they demonstrate the crucial need to consider both IOP and ICP to fully understand pressure effects and likely susceptibility to glaucoma. Our results highlight the complexity of LC biomechanics, and the challenge to understand the multiple interacting factors. We expand upon these points below.

### Changes in either IOP or ICP caused significant, detectable ONH deformations

IOP and ICP were both manipulated, allowing us to detect effects of IOP and ICP individually. Whereas many studies have described effects on the ONH of variations in IOP ^10,35–40^, far less understood are the effects of modulating ICP and its interactions with IOP. Our results demonstrate that the effects of ICP on the ONH can be detected in vivo, and that they can have a magnitude comparable to the effects of IOP.

ONH structures were manually delineated at relatively high density compared with previous studies ^41–43^. For instance, we marked the ALC every 31 μm, about six times higher than previously ^43^. The dense markings allowed us to reconstruct detailed BMO planes and ALC surfaces, which were then used to quantify their changes and deformations in 3D. A major concern of ONH studies based on OCT data is the variable visibility, particularly that of the deep structures like the LC ^26,44–46^. Inconsistent visibility means that a simple comparison of parameters across conditions has a substantial risk of being biased by region visibility. To avert this bias, all the LC analyses in this work were based on uniform sampling of regions that were clearly visible in all pressure conditions of an eye. This is similar to the shared sectors used by Strouthidis et al ^31^ and the overlap restriction in Wang et. al. ^21^. This approach reduced the size of the lamina regions analyzed from an average of 57% to 34%, which still compares well with those of other studies ^21,31^. In contrast with previous work which considered deformations of ONH microstructure ^21^ this study places focus on large-scale parameters. This allowed us to observe changes in the overall size and shape of the scleral canal.

Acute modulation of ICP and IOP allowed us to measure the deformations without potential confounding from remodeling or inflammation associated with chronic pressure changes ^47,48^. Another strength of this work is that we studied monkeys, the animal model that most resembles humans ^49^. The procedures we performed were highly invasive and pose multiple risks in human subjects. Recently, Fazio et al have overcome many of these by studying IOP and CSFP in brain-dead human organ donors.^27^ Fazio et al ^27^ reported on ONH deformations in response to changes in IOP and CSFP in brain-dead human organ donors. In this study, ICP was estimated based on donor body position and not probed or controlled directly as done here. Our in vivo measurements also avoid postmortem effects, such as tissue processing, histology and the absence of blood pressure ^26^.

### There were strong interactions between the effects of IOP and ICP on the ONH

Here, we presented robust evidence that the effects of ICP on the ONH can be influenced by the level of IOP, and conversely that the effects of IOP can be influenced by the level of ICP. Interaction of ICP and IOP-mediated effects were observed in all eyes and in the majority of parameters. This demonstrates the importance of considering the effects of both ICP and IOP together rather than only independently. When possible, incorporation of ICP as an experimental variable in studies of IOP-induced deformation is warranted.

An important consequence of the complex interactions between IOP and ICP was that **elevating both pressures by the same amount, and thus maintaining TLPD constant, did not cancel out the pressure effects on ONH visibility and morphology**. We found that none of the tested eyes had the same structural measurements when subject to the same TLPD but different IOP/ICP conditions. A constant translaminar pressure did not ensure constant ONH morphology. A better understanding of how these interactions are influenced by both micro- and macro-scale ONH structure can allow us to determine the factors necessary to better predict these complex structural responses and how they might impact the health of the resident neural tissue. This indicates that TLPD is unlikely to be a strong parameter to predict morphologic changes in the LC, as has been proposed ^50^.

### A high degree of heterogeneity in responses to pressure manipulation was observed among eyes

With variations in either IOP or ICP, no parameters changed consistently in one direction for all eyes. This heterogeneity cannot be explained by variability in marking or measuring of ONH structures, as demonstrated by the high repeatability of measurements, the narrow confidence intervals for many parameters, and consistent findings in the bootstrap analysis. The bootstrap analysis increased confidence of the observed gross structural changes. Out of all experimental configurations considered in this study, ~70% (44/65) of structural changes in response to elevations in either ICP or IOP were significant. This indicates that we observed robust changes following pressure manipulation. Furthermore, the absolute observed percentage changes were as large as 123%, 15%, 53%, and 33% for canal planarity, canal area, LC depth, and LC visibility, respectively. Conversely, changes in canal aspect ratio were relatively small: within 4%. It is worth reminding the reader that the experiment and analysis were not designed to determine whether the responses to IOP and ICP variations measured in these monkey eyes will extend to other eyes or monkeys. This cannot be determined from the small set of eyes studied. The bootstrap analysis shows that the deformations measured are likely to be ‘true’ changes in the structures as visible in the OCT scans, and not due to statistical noise or variability in the markings. Additional studies with more eyes and animals are necessary to characterize the population and the variable directionality of the observed tissue responses.

Our previous study on the response of microstructure to IOP and ICP modulation showed that the best fit statistical models included an interaction between ICP and TLPD ^21^. In that study, there was similarly a marked variability in responses between eyes. Both the prior and current studies emphasize the importance of considering both IOP and ICP in evaluation of the ONH. The current study, focused on macro-scale ONH features, suggests that the variable micro-scale responses previously observed extend to global measures of deformation. This may help to explain the variable responses of patients’ eyes to elevated pressure in certain disease states (i.e. glaucoma, intracranial hypertension). These findings also suggest that a more personalized medicine approach to optic neuropathy may be optimal for determining the risk and best course of treatment for individual patients. Further work is necessary to understand how ONH structural factors are associated with increased glaucoma risk.

A few studies have characterized the effects of acute manipulations of IOP and ICP on prelaminar neural tissue displacement. Zhao et al. showed posterior movement of the ONH surface and surrounding peripapillary retina with IOP elevation, and greater displacement at lower ICP using a rat model ^18^. These results are concordant with the study conducted by Morgan et al. with dogs showing posterior displacement of the disc surface with IOP elevation, whereas CSFP elevation prompted anterior displacement. This study also reported non-linear surface deformations as a function of TLPD where most displacement occurred in the low range translaminar pressure gradients ^15^. In our 2017 study of 5 monkeys ^21^, we similarly modulated IOP and ICP, stepwise, at a range of pressure settings. We observed a significant interaction between the effects of IOP and ICP on changes in LC microstructure. It is possible that the model used for the study described here also exhibits similar non-linear behavior. A comprehensive characterization of the deformations in the monkey ONH at more pressure combinations, as done previously in dogs^15,51^, is necessary to determine this.

Feola et al. ^14^ utilized phase-contrast micro-CT to capture CSFP-induced deformations of the LC and retrolaminar neural tissue in an ex vivo porcine eye model. They found that variation of cerebrospinal fluid pressure greatly impacted the distribution of strain within the RLNT and to a lesser degree, the LC as well. In line with the heterogeneity of the observations reported here, the spatial distribution of strains within the LC differed greatly among individual eyes. Understanding the factors that contribute to this heterogeneity of responses would be of great value in prediction of medical risk. Numerical models are shedding valuable light in the mechanistic interactions between the forces acting on the ONH, including IOP, ICP, blood pressure and tension from the optic nerve ^20,52,53^.

Epidemiologic work has reported a correspondence between TLPD and both structural and functional glaucomatous changes ^13,54–56^. In human subjects, the Valsalva maneuver was similarly shown to cause a greater acute increase in cerebrospinal fluid pressure than IOP, resulting in changes of ONH morphology ^50^. Transiently altered TLPD was associated with decreased cup/disc ratio as well as maximum optic cup depth. Many of these studies involve subjects with chronic elevation or suppression in either ICP or IOP. As different IOP/ICP combinations with the same TLPD did not result in consistent deformation of the ONH, our results suggest that with acute manipulation, the interaction may be more involved. This is consistent with the findings of the numerical studies mentioned above. For instance, Hua et al found that the overall influence of TLPD was 28 times smaller than that of IOP, and weaker even than CSFP whose effect were 16 times smaller than those of IOP. In contrast, it is also important to note that there are a number of excellent papers which do discuss IOP/ICP in more detail. They do not trivialize TLPD and instead argue for the importance of dealing with IOP and ICP in a more nuanced way ^20,52,57,58^. Our results support this conclusion as well.

In addition to the strengths, the limitations of this study must be recognized to best inform future directions. To evaluate the ONH response to acute, controlled changes in IOP and ICP, we focused on two structures with well-established relevance in analyzing effects of IOP, the ALC and the scleral canal ^9,11,59^. Future studies will need to incorporate analysis of additional factors known to influence mechanical insult in the ONH, such as cerebrospinal fluid pressure ^14^ and additional ONH structures, such as blood vessels ^60^ or even cellular components ^61–63^. This can allow us to better understand the full effects of the pressure-induced changes. Besides macroscopic deformations of the scleral canal and ALC that were investigated in our study, microarchitecture, such collagen crimp ^64^, could be an important measurement as well. Analysis of these factors will help to better explain the reason behind and the implications of macroscopic deformations reported here. Additionally, this will aid in assessment of whether particular regions of ONH could be loaded with more or less force in response to changes in IOP and/or ICP, and how this may eventually contribute to neural tissue insult ^65^. We accounted for transverse scaling by relating OCT scans to histology in order to ensure an accurate baseline scale. Changes in axial length due to IOP changes could potentially result in scaling differences that were not accounted for. Lastly, the experimental protocol included setting several more IOP and ICP conditions than what we analyzed in this work. We decided to select a subset of pressures as a first analysis to evaluate the effects and potential interactions between IOP and ICP. Future studies should use a more comprehensive set of the data acquired. It is important to acknowledge the additional IOP and ICP conditions between the ones studied. These steps are important because they allow the eyes to stabilize after pressure changes, ensuring that our measurements are free from viscoelastic effects.^21,24,26,27^

We focused on the effects of acute variations in ICP and IOP. It is likely that the effects of chronic exposure to variable levels of ICP and/or IOP will have effects that are different from the acute ones, such as remodeling and inflammation. Understanding the role of ICP and IOP in glaucoma will require a careful characterization of the chronic effects of these pressures. We posit that understanding the acute effects of the pressures, as advanced in this work, is an essential and necessary step. In other words, to understand the long term process of glaucoma, we must also understand the short term biomechanics of the ONH. An improved understanding of acute and short term interactions between ICP and IOP may also be relevant in the development and screening of techniques to measure ICP non-invasively. Some techniques, for instance, are based on the concept of TLPD, which our study suggests may be problematic.^14,66^

A technical challenge for our analysis was the lack of an absolute frame of reference. This is a limitation that our study shares with other work on ONH morphometrics ^67^ Although the BMO plane has been commonly used as a reference for measurements within the ONH ^38,68,69^, pressure changes and pathology can cause BMO surface deformations. In this work, the effects of these deformations were minimized by using BMO best-fit plane. For this reason, analysis between pressure settings and structural registration was based on the centroid and principal axes of the BMO best-fit plane. We chose this method because it is objective and repeatable, facilitating inter and intra-study comparisons. However, as we and others ^70^ have shown, the BMO itself is affected by IOP and ICP. Hence, it is possible that changing the registration would produce slightly different results. Future work should consider other potential methods of registration and measures of the ONH that are independent of the BMO ^27,71,72^.

Because the subarachnoid space of both eyes and the brain are directly connected, it is not possible to manipulate ICP independently between two eyes within the same animal. The effects of ICP could be more profound in an eye that had been exposed to elevated ICP for a longer period of time. This would be the case in monkeys where both eyes were imaged, such as M3. One eye of M3 was exposed to ICP elevation prior to imaging due to earlier imaging of the contralateral eye. Interestingly, deformations of the scleral canal and ALC were larger in M3L when compared those in M3R. This was particularly profound in the cases of canal planarity and ALC depth.

It is important to articulate not only which changes were statistically significant, but which may be physiologically impactful. With limited information available about the risks associated with these particular degrees of ONH deformation on tissue health and vulnerability, it is not yet straightforward to determine which of these factors are associated with medically relevant risk. From a biomechanical perspective, studies suggest that it is often not the magnitude of the displacements, but their gradient (deformations) that are best predictors of damage.^73,74^ Answering these questions will require separate investigations in future work.

We manipulated and measured ICP at the brain, whereas it is the cerebrospinal fluid pressure immediately behind the globe that directly impacts the ONH. ^12,13,15,51,54^ Although these two pressures are closely related, they are not necessarily identical, with likely differences in their magnitude and potentially even a time lag between changes in ICP translating to pressures within the orbit. It is still unknown if the 5 minutes we waited before imaging after a change in pressure are sufficient to allow the changes in ICP to fully translate to the orbit.

In summary, our study provided evidence of substantial changes in gross ONH morphology caused by acute changes in IOP and ICP that were unique to each individual eye. Additionally, we describe a significant interaction between the effects of ICP and IOP on the ONH scleral canal and LC. Altogether, our results show that ICP affects sensitivity to IOP, and thus that it can potentially also affect susceptibility to glaucoma.

## Acknowledgments

We thank Huong Tran for help throughout the project.

## Notes

### Competing Interest Statement

Proprietary Interest: J.S. Schuman receives royalties for intellectual property licensed by Massachusetts Institute of Technology to Zeiss. All other authors: Nothing to disclose

### Summary of Updates

Comprehensive revision from abstract to discussion. New figures.

