## Supplemental Figure 1 for "Interplay between intraocular and intracranial pressure effects on the optic nerve head in vivo"

### Supplemental material

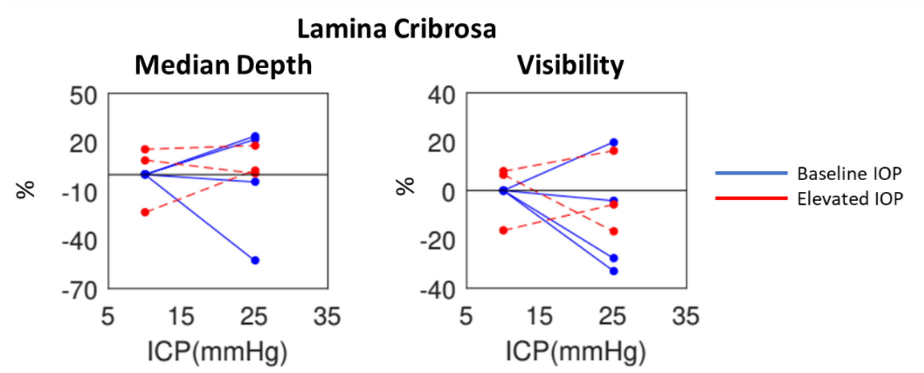


**Supplemental Figure 1:** Percentage changes of lamina median depth and lamina visibility with respect to baseline values due to ICP elevation at baseline IOP (blue) and elevated IOP (red). In some cases, changes in ICP have substantial effects on both lamina depth and visibility, often quite different depending on the level of IOP. Each line represents the estimate for one eye.
